# “In vitro construction and long read sequencing analysis of a 24 kb long artificial DNA sequence encoding the Universal Declaration of the Rights of Man and of the Citizen ”

**DOI:** 10.1101/2023.06.26.546242

**Authors:** Julien Leblanc, Olivier Boulle, Emeline Roux, Jacques Nicolas, Dominique Lavenier, Yann Audic

## Abstract

In absence of DNA template, the *ab initio* production of long double-stranded DNA molecules of predefined sequences is particularly challenging. The DNA synthesis step remains a bottleneck for many applications such as functional assessment of ancestral genes, analysis of alternative splicing or DNA-based data storage. We propose in this report a fully *in vitro* protocol to generate very long double-stranded DNA molecule starting from commercially available short DNA blocks in less than 3 days. This innovative application of Golden Gate assembly allowed us to streamline the assembly process to produce a 24 kb long DNA molecule storing part of the Universal Declaration of Human rights and citizens. The DNA molecule produced can be readily cloned into suitable host/vector system for amplification and selection.

## Introduction

For billions of years, double-stranded DNA molecules (dsDNA) have been the molecular support of choice for the storage of biological information and the support of life. In biological systems, DNA replication is a core biological process that occurs at high speed in prokaryotes (∼ 700 nucleotide/sec, ^1^). In eukaryotes, the process is slower (∼ 15-30 nt/sec ^2^) but so highly parallelized that it allows to achieve replication of a 1,7 billion nt genome in less than 30 min in early Xenopus embryos (∼ 9.4 × 10^5^ nt/sec). It is also particularly efficient for the synthesis of collinear DNA molecules of up to hundreds millions of nucleotides such as lungfish chromosomes (^3^). This is made possible by the faithful copy of pre-existing nucleic acid polymers and template-specific DNA polymerases (^4^). On the other hand, the *ab initio* production of DNA molecules of only thousands of nucleotides of defined sequence remains a technological and scientific challenge. So far, commercial companies generally offer cost-effective chemical synthesis of oligonucleotides up to a few tens of nucleotides and advertise longer DNA molecules of desired sequences for up to 50,000 nt (^5,6^) albeit at a substantial cost (**Table 1**). Assemblies of impressively long DNA molecules such as the mouse mitochondrial genome (16.7 kb), T7 bacteriophage (39.9 kb) or even the *Mycoplasma genitalium* genome (583.0 kb) are documented in the literature but all rely on hierarchical assembly and molecular cloning steps which are time-consuming and laborious (^7–9^). Furthermore, analysis and assessment of the accuracy of the assemblies relies on their cloning into an appropriate vector, which must be transferred into a host and biologically selected. A less laborious procedure has yet to be defined, aimed at producing *in vitro* and cost-effectively long dsDNA molecules of predefined sequences in the absence of a DNA template.

**Table 1.**
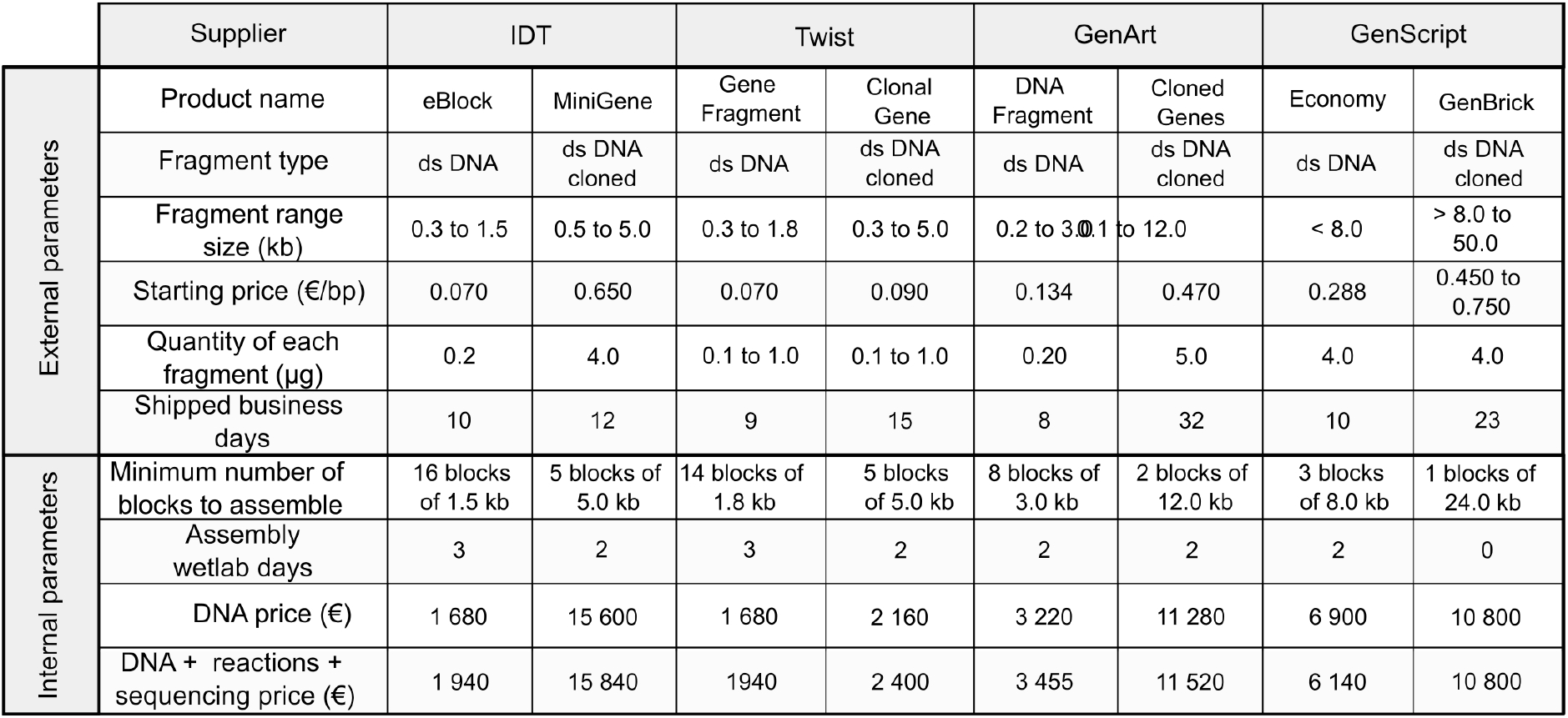
Comparaison of production costs for the construction of one dsDNA fragment of 24 kb using DNA parts from differents suppliers. (2023 prices). Initial DNA parts are either obtained as linear DNA molecules or as DNA molecules cloned into a vector.

This limitation is unfortunate considering the numerous applications of long DNA molecules in synthetic biology (^10,11^). Long synthetic DNA molecules are also a subject of interest in experimental biology, as they can provide insights into several areas of research such as the reconstruction of ancestral DNA genes, regulatory elements and proteins (^12^) or the investigation of alternative splicing regulation (^13^). Moreover, the production of large artificial or semi-artificial DNA molecules, including mutants or variants, is a widespread practice for the remodelling of genetic circuits, the evaluation of gene function, and the identification of functional domains within DNA or encoded proteins (^6^). Additionally, these DNA molecules are also useful as templates for RNA-vaccine production (^14^). Their main industrial applications concern *in vitro* screening of mutations for the development of therapeutics and chemical products, including drugs and biofuels (^10,15^).

Another important use of long DNA Molecules takes root in their dense and stable information content. Current data storage systems, whether magnetic or silicon-based, are indeed unable to respond to the rapid increase in archiving needs and have several drawbacks such as a limited lifespan, energy consumption, miniaturization restrictions and environmental impact. DNA molecules are nowadays envisaged as an alternative to store data because of their tremendous information density and high chemical stability as evidenced by the recovery and analysis of ancient DNA extracted from fossils (^16,17^). However, the main bottleneck for the storage of information on DNA is the DNA synthesis step itself, which is slow, expensive and can only generate short oligonucleotides (^18,19^). The reduced size of the oligonucleotides implies fragmenting the digital documents into a large number of small pieces which must necessarily include indexes allowing the reconstruction of the original document (^20^). The size of this index can be important compared to the size of a DNA oligonucleotide and thus significantly affect the effective amount of information stored and the synthesis costs. Incidentally, this fragmentation will also increase the difficulty of recovering the original documents. On the other hand, single molecules real-time sequencing (Pacific Bioscience) or Nanopore sequencing (Oxford Nanopore Technology) are now capable of reading individual long DNA molecules, which is compatible with storing of information on longer DNA fragments (^21–23^). Such long DNA molecules will only contain a small fraction of indexing information making them more archival-efficient.

Many methods have been developed to construct large DNA fragments from short chemically synthesized DNA oligonucleotides (^6,15,24^). The most common are Golden Gate assembly (GGA), Gibson cloning and polymerase cycling assembly. These methods are all based on user-defined overlapping ends, defined as overhangs, and generally allow the seamless joining of 5 to 10 fragments per reaction, the assembly is then cloned into a vector, transferred to an host, selected and amplified.

Herein we propose a fully *in vitro* iterative method based on GGA to faithfully construct a long synthetic DNA molecule starting from commercially available dsDNA. The synthetic DNA molecules could be used for any common biological purpose or to store information. The various assembly steps of our procedure were evaluated on an application for storing part of the Declaration of the Rights of Man and of the Citizen (26th August 1789) in a single long DNA fragment which was then sequenced using Oxford Nanopore Technology (ONT) to retrieve the original text. The different steps of the DNA construction were analysed using ONT long read sequencing technology to quantify the ordering of the building blocks, demonstrating that our protocol is highly effective in obtaining correctly assembled molecules.

## Results and discussion

Articles 1 through 9 of the Declaration of Human and Citizen rights were encoded into a 23.4 kb long DNA sequence (**Supplementary data 1**). To fully build *in vitro* this dsDNA molecule we started from 50 commercially available 524 nt long dsDNA molecules (eBlocks, IDT™) that we assembled into 4,764 bp long dsDNA molecules (BigBlocks) that were then assembled into a 23,796 bp DNA molecule (MaxiBlock) (**Figure 1**). This fully *in vitro* strategy uses as building blocks commercial dsDNA molecules few hundreds of nucleotides long (eBlock, IDT™). The design of these building blocks relies on an architecture composed of, starting from the end, a 15 nt long buffer sequences that ensure the integrity of the two *BsaI* prefix and suffix sequences, the four bases cleavage sites that allow directional ligation and finally the cargo part of the DNA containing the encoded information (**Figure 1A**). Upon cleavage by *BsaI*, the DNA cargo part will remain solely framed by the predefined 4 nt overhangs to allow ordered assembly of 10 blocks in a single GGA reaction. Specific primer pairs targeting the extremities of the assembled BigBlocks (**Figure 1B**) are used to select and amplify the 4,764 bp long assembly products and flank them with sequence specific type IIS (*BsaI*) restriction sites to allow for the second *in vitro* assembly, to generate the MaxiBlocks (**Figure 1C**). As described in **Supplementary data 2**, the different assembly steps were controlled during the course of the experiment. The 23,796 bp MaxiBlocks are PCR amplified before utilization.

**Figure 1.**
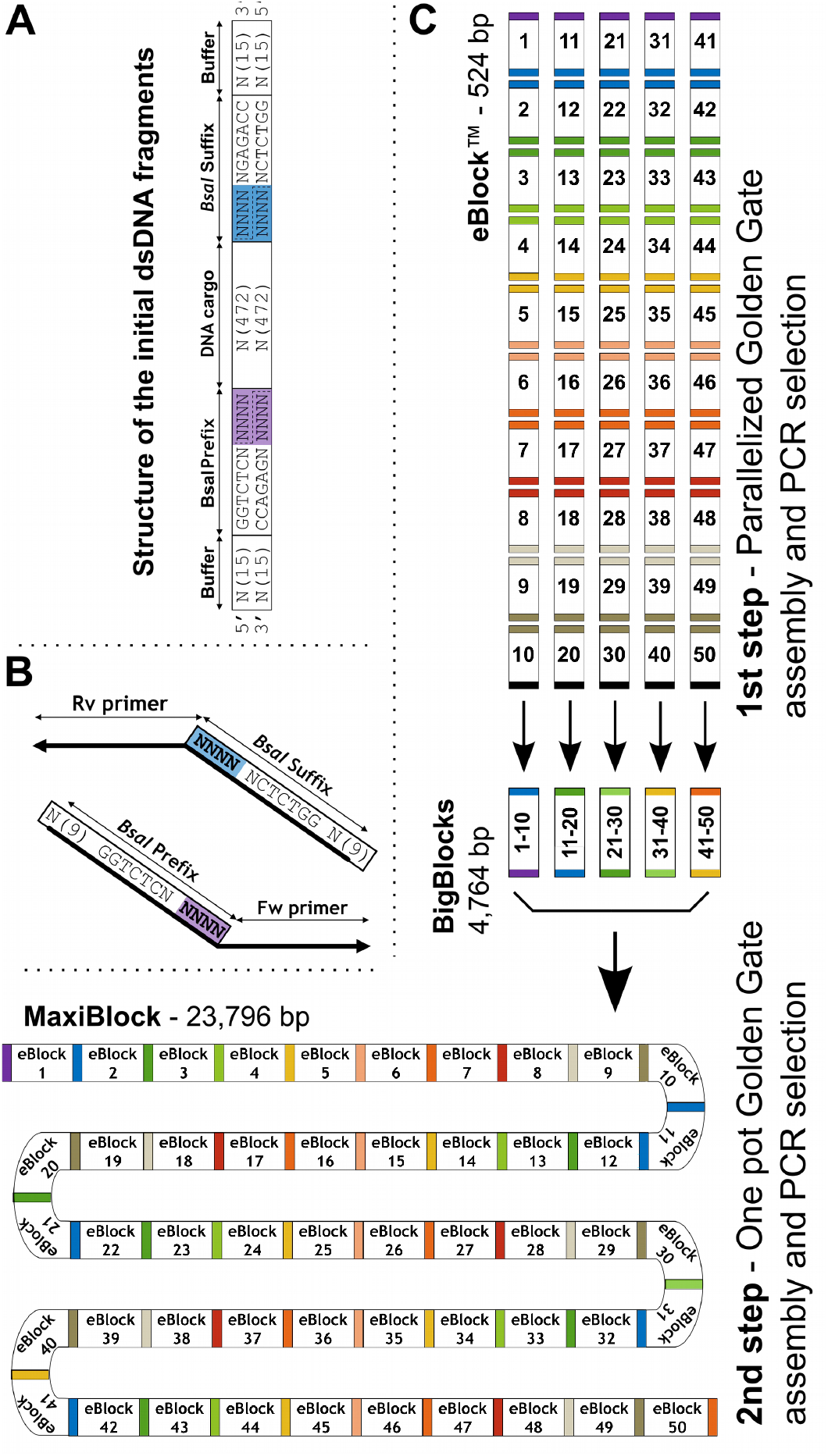
Overview of the strategy for building 23,796 bp dsDNAfrom short dsDNA fragments. **A)** dsDNA building blocks are composed of a DNA cargo framed by 2 *Bsal* sites and a buffer sequence at the extremities. The colored tetranucleotides correspond to the overhang produced by the Bsal digestion during assembly. **B)** Oligonucleotides are composed from 5’ to 3’ of a 9 nt buffer sequence, a *Bsal* site and a BigBlock specific sequence. **C)** Two steps strategy for building a 23,796 bp dsDNA. 5 sets of 10 eblocks are assembled into Bigblocks (4,764 bp) in parallel using GGA. The expected products are selected, amplified and decorated with *Bsal* sites using specific primer pairs described in B. Bigblocks are mixed in equimolar amounts before proceeding to GGA. The final 23,796 bp product is selected and amplified by PCR.

### First assembly step: eBlocks to BigBlock

To design a 23,796 bp long DNA molecule, we split the sequence into fifty 524 bp parts, which were first assembled in groups of ten to generate five 4,764 bp BigBlocks. At the time of our study, purchasing the 524 bp eBlocks at Integrated DNA Technologies (IDT™) was the most cost-effective way to obtain these long dsDNA molecules (**Table 1**). The five GGA reactions aiming to obtain 4,764 bp long DNA molecules were directly controlled on agarose gels (**Figure 2A**). Incomplete sequential ligation occurred and a scale of DNA molecules ranging from 480 bp, the initial size of the eBlocks after *BsaI* digestion, to 4,764 bp by incremental steps of about 500 bp could clearly be observed. We selected the desired 4,764 bp assembly using a PCR targeting the extremities of the BigBlocks. As presented in **Figure 2B**, size fractionation of the PCR product demonstrated that for each of the five BigBlocks we could specifically amplify the longer 4,764 bp DNA molecules. This demonstrated that ∼ 5 kb long dsDNA molecules can be effectively assembled *in vitro* from commercial 524 bp dsDNA molecules. However so far this did not demonstrate that the fragments are correctly ordered in the assembly.

**Figure 2.**
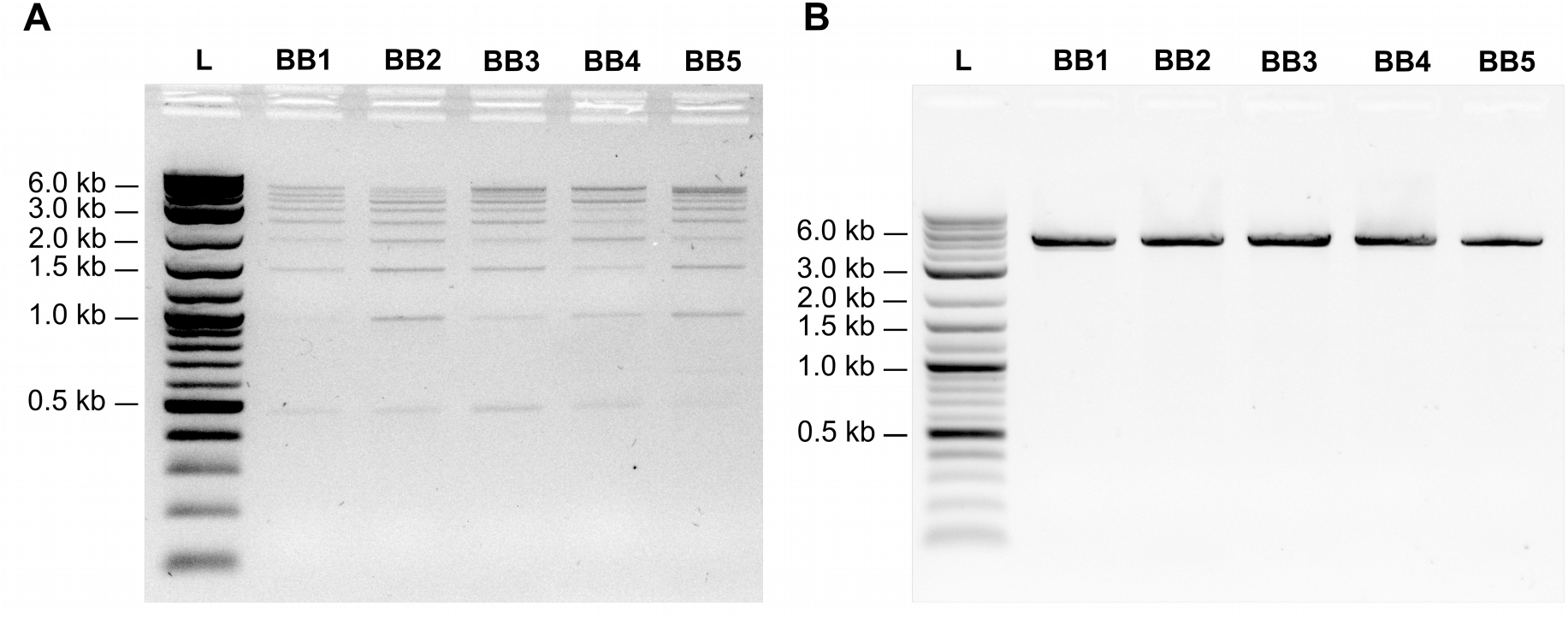
BigBlocks assemblies before and after PCR analyzed on 2% agarose electrophoresis gels. **A)** Five different BigBlocks (BB1 to BB5) are assemblies from 50 eBlocks of 524 bp (IDT eBlocks™). Each BigBlock is composed of 10 oriented eBlocks. Expected size = 4,764 bp. **B)** BigBlocks products from (A) are PCR-amplified with dedicated flanked primers. Expected size = 4,796 bp. (L) indicates the 1 kb Plus DNA ladder (NEB).

To determine to which extent the assembly process is ordered, we took advantage of ONT sequencing technology to directly sequence ligation and PCR products for BigBlock 1 and 4. We reasoned that single molecule long read sequencing technology such as ONT or PacBio were the most effective in obtaining sequence information about the accuracy of long collinear assembly. For the assembly of BigBlock 1 (**Figure 3A)** or BigBlock 4 (**Figure 3B**) the graphics represent the percentage of transition from block (n) to another block, to the end of the read (end), or to an unrecognized sequence (Unknown). BigBlock 1 graphics were produced from 4,935 reads with median length of 1,435 nt and median quality of 10.9. For BigBlock 4, graphics were generated from 5,281 reads with median length of 1,898 nt and median quality of 11.1 (**Supp data 3**). These medians reads lengths are about one-third of the expected product size in agreement with size fractionation (**Figure 2**), showing that the reaction is composed of partially assembled molecules. Directly analysing the assembly reaction products, we could note that the percentage of correct concatenations of two consecutive eBlocks ranged from 61 to 95% for the assembly of BigBlock 1 and from 74 to 94% for the assembly of BigBlock 4 (Left panel on **Figure 3 A and B** respectively). In both cases and as expected, eBlock 10 was most generally terminal, 99 and 96% respectively. In the case of BigBlock 1 assembly, the lower efficiency of assembly of eBlock 5 was associated with either incorrect assembly of eBlock 5 to eBlock 7 (∼ 10%) or eBlock 5 being terminal (∼ 20%). Similarly, in the assembly of BigBlock 4, the lower level of concatenation of eBlock 34 to eBlock 35 was due to eBlock 34 being terminal. This analysis indicates a faithful assembly process as most generally the assembly is correct from eBlock (n) to eBlock (n+1), or otherwise no assembly occurs.

**Figure 3.**
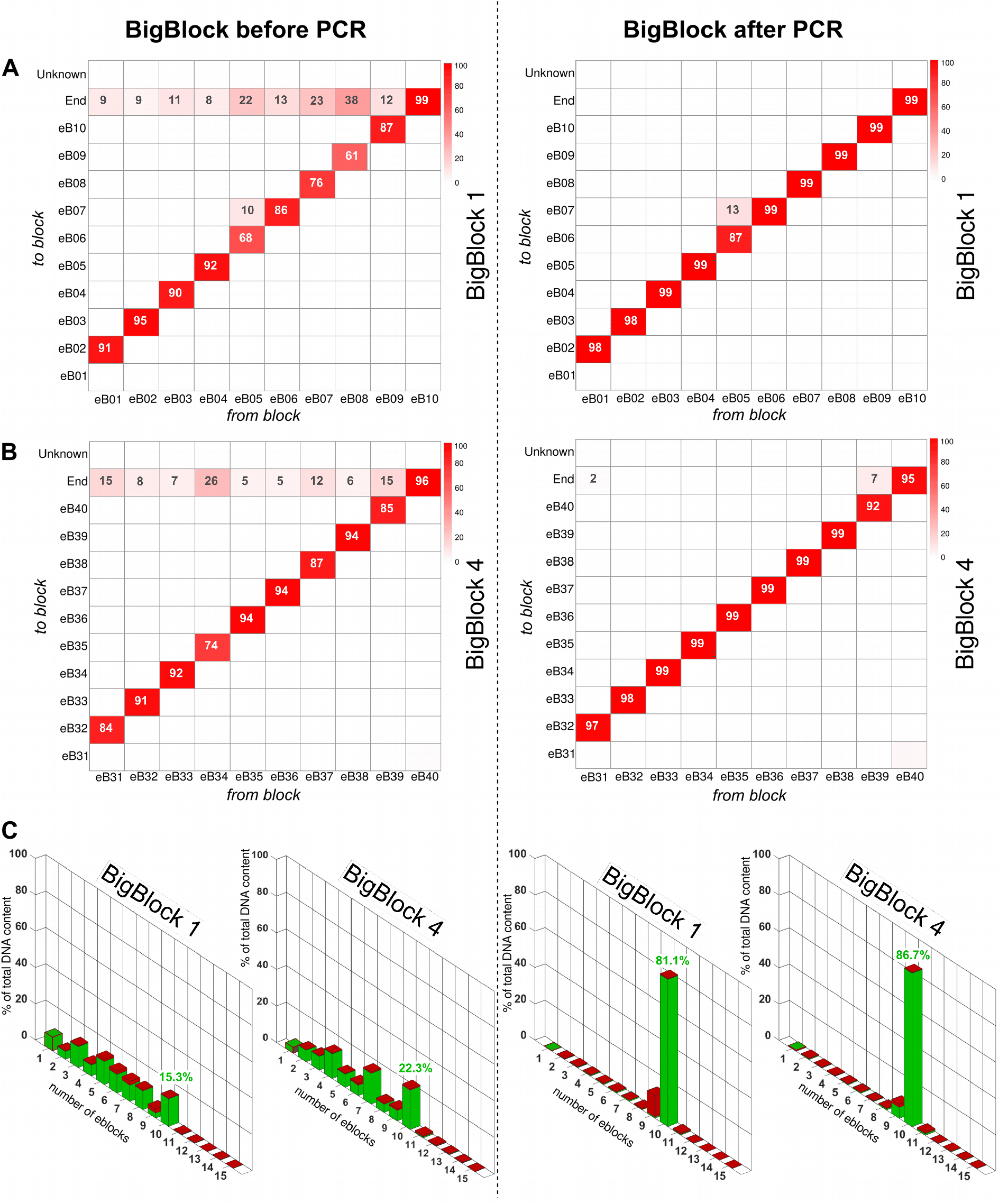
Analysis of the assembly of BigBlock 1 and 4 before (left panel) and after (right panel) PCR selection. **A)** Heatmap presenting in the assembly of BigBlock 1 the proportion (%, only percentage above 1% are shown) of two ligated consecutive eBlocks determined by ONT sequencing. **B)** Same as **A** for BigBlock 4. Sequencing data are from assemblies shown in Figure 2. Unknown, eBlock is ligated to an unknown DNA sequence. End, the eBlock is terminal. C) Distribution of the DNA content in the assembly for BigBlock 1 and BigBlock 4 relatively to the number of eBlock composing the reads. Green, all the eBlocks composing a read are in correct order. Red, at least one eBock is misplaced.

We used PCR to select and amplify the full-length BigBlock assembly (see **Figure 2B**). BigBlock 1 post-PCR graphics were produced from 3,000 reads with median reads length of 4,757 nt and median read quality of 10.9. For BigBlock 4 post-PCR, graphics were generated from 1,476 reads with median length of 4,752 nt and median quality of 10.9 (**Supp data 3**). PCR selection and amplification therefore led to a ∼ 3-fold increase in median reads length compared to pre-PCR products, 4,757 nt compared to 1,435 nt for BigBlock 1 and 4,752 nt compared to 1,898 nt for BigBlock 4. This median size is close to the 4,796 nt size of the expected product. This illustrates how effective is the PCR amplification step in selecting full length assembly products.

After PCR, the BigBlock sequencing results show a frequency of correctly ordered eBlock pairs ranging from 87-99% for BigBlock 1 assembly and 92-100% for BigBlock 4 assembly, a neat increase in accuracy with respect to previous results (Right panel on **Figure 3 A and B** respectively). The problematic assembly of eBlock 5 to eBlock 7 in BigBlock 1 before PCR remains at a similar level after, while the premature termination of the assembly at eBlock 5 becomes negligible as it cannot be amplified. Similarly, for BigBlock 4, only the premature termination of the assembly at eBlock 9 remains notable (7%), while being reduced by half compared to the assembly without PCR. This shows that when an eBlock is erroneously terminal this does not affect the final PCR-selected product, while if two eBlocks are erroneously assembled they will stay present in the PCR-selected assembly. It is therefore important to maximize individual ligation efficiency.

While the previous analysis focuses on the correct assembly of pairs of eBlocks, it is also important to estimate the overall accuracy of whole BigBlocks assembly. We used the above ONT sequencing data to estimate the distribution of the length of the DNA assembly and to determine the proportion of correct assembly for the different fragment sizes. This also enabled us to quantify the enrichment in correct assemblies following the PCR amplification and selection process (**Figure 3C**). As observed by gel electrophoresis, the reaction generated all product sizes between 1 and 10 assembled eBlocks (0.5 to 5 kb). The target molecule composed of 10 eBlocks dominates the reaction products and represent 15 and 22% of the sequenced reads in BigBlock 1 and BigBlock 4 reactions respectively with less than 3% inaccurate assemblies. The proportion of molecules with more than 10 eBlocks is negligible (< 1%). Upon PCR amplification and selection, we have observed a drastic depletion of BigBlocks composed of less than 10 fragments. About 84% of the DNA is included in molecules composed of 10 fragments almost all in the right (∼ 99% for both BigBlock 1 and BigBlock 4).

Taken together these analyses indicate that the combination of directed assembly without any intermediate purification step and with PCR selection allows to efficiently produce long DNA molecules of the expected structure.

### Second assembly step: BigBlocks to MaxiBlock

Having established the reliable production of 4,796 nt dsDNA molecules (the BigBlocks), we set about assembling five of them into a 23,796 bp MaxiBlock. As shown above, the five BigBlocks were amplified using primers that introduced *BsaI* restriction sites together with tetramer sequences required to correctly order the fragments (see **Figure 1B)**. The assembly reaction of the five BigBlocks shown in **Figure 2** was performed using GGA. The reaction products were analyzed either directly after the reaction or after PCR amplification and selection using primers targeting the first (BigBlock 1) and the last (BigBlock 5) BigBlocks of the assembly. Because of the expected size of the DNA assembly, we analyzed the reaction products by capillary electrophoresis on a Tape Station (Agilent Technologies) (**Figure 4**). It is noteworthy that the resolution of the Tape Station does not allow for precise sizing of the DNA molecules in the higher size range. The products of the ligation reactions (**Figure 4A, MB1**) ranged in size from 4 kb to more than 15 kb in a manner compatible with partial and complete assembled molecules in the reaction mix. After PCR amplification (**Figure 4B, MB2**) using primers targeting the extremity of the Maxiblock and LongAmp^®^ Hot Start Taq (NEB^®^) DNA polymerase, a single PCR products of size between 15 and 48 kb is detected and in agreement with the expected 24 kb DNA product.

To better evaluate the quality of the assembled fragments we again took advantage of ONT sequencing to compare the reaction products before and after amplification and selection of a 23,796 bp DNA assembly. ONT sequencing of MaxiBlock assembly produced 25,507 reads with median length of 3,091 nt and median quality of 11.7 before PCR and 3,215 reads with median length of 1,143 nt and median read quality of 11.9 after PCR (**Supp data 3**). The limited median size of the sequencing reads of the MaxiBlock assembly takes its root in the presence of a) numerous short sequencing reads composed of part of terminal eBlocks 01 or 50 and b) presence of sequences that were ill-attributed during demultiplexing **(Supp data 4)**. As shown in **Figure 5 A, left panel**, the proportion of correctly ordered BigBlock pairs ranges from 56 to 85%. As designed, the BigBlock 5 is most often (97%) the terminal block. When BigBlock (n) is not ligated with BigBlock (n+1), then it is usually a terminal block of the assembly. Upon PCR selection the DNA molecules sequenced are mainly composed of an assembly of BigBlocks in the right order (**Figure 5B, right panel)**. The proportion of correctly ordered BigBlock pairs ranges from 89 to 96% and BigBlock 5 is exclusively terminal. This indicates that upon PCR, full-length products are selected while a large proportion of partial assemblies are not amplified.

**Figure 5 :**
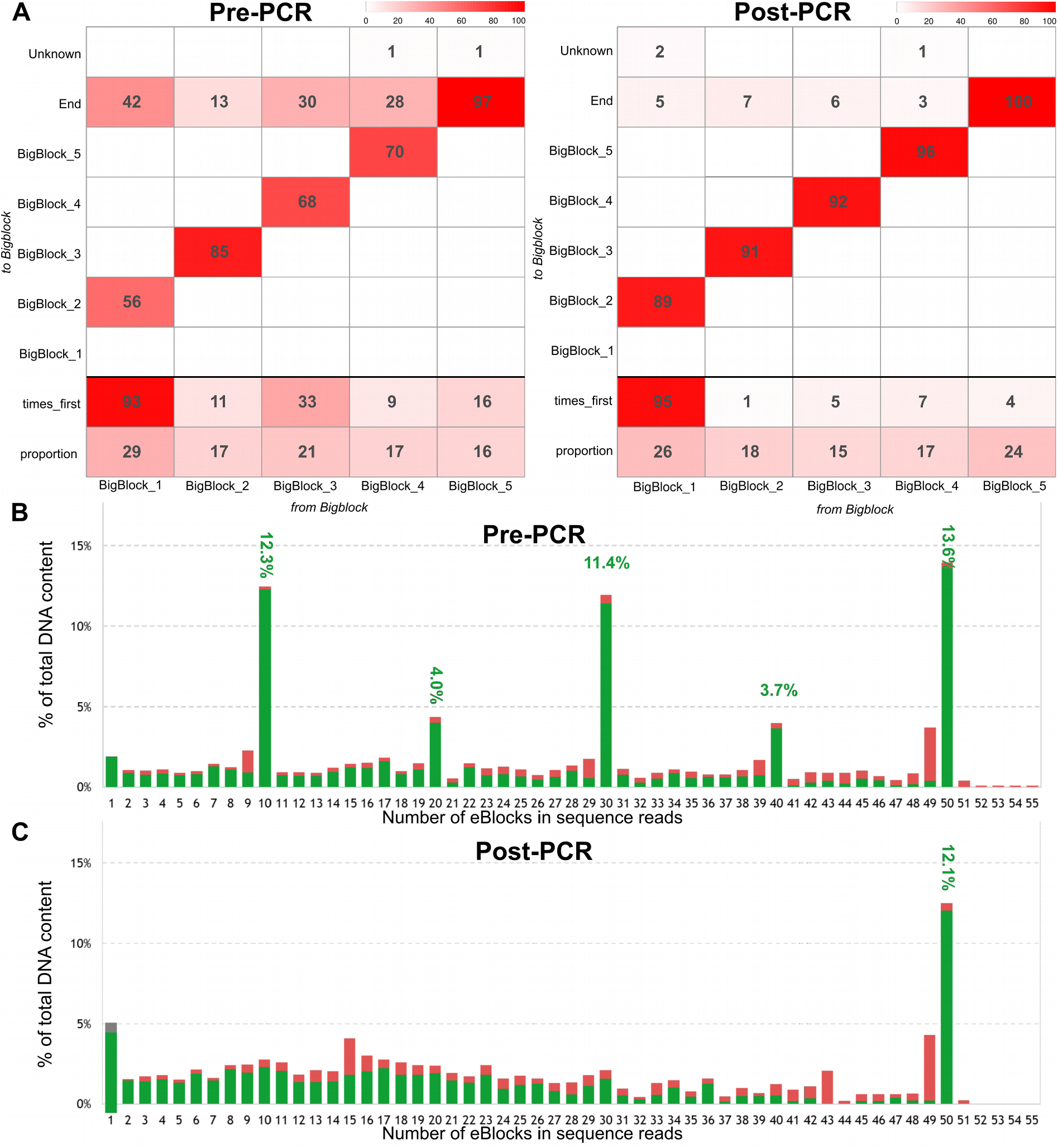
ONT sequencing analysis of MaxiBlock assembly. **A)** Heatmap showing the proportion of pairs of linked BigBlocks, « from block on the x-axis, « to » block on the y-axis (values >1% are shown) before and after PCR selection. At the bottom of the table appear the overall proportion of each BigBlock in the set of BigBlocks appearing in the Maxiblocks (row « proportion », sums to 1), and the fraction of Maxiblock containing this BigBlock in first position (row « times-first », relative to the proportion below). Sequencing data are from assemblies shown in Figure 4. Unknown, BigBlock ligated to an unknown DNA sequence. End, BigBlock is terminal. B) Distribution of the DNA content in the MaxiBlock before PCR selection assembly relatively to the number of eBlocks composing the reads. C) Same as B) after PCR selection. Green, all the eBlocks composing a read are in correct order. Red, at least one eBlock is misplaced.

We analyzed the number of eBlocks and the correctness of their assembly for all sequenced reads. In the pre-PCR reaction products, a striking pattern of reads composed of 10, 20, 30, 40 and 50 eBlocks are present and represent the assembly of 1, 2, 3, 4 or 5 BigBlocks together. They respectively represent 12, 4, 11, 4 and 14% of the DNA content of the reaction products. Most notably the majority of these reads are composed of correctly ordered fragments (**Figure 5B**).

As done for the selection of the BigBlocks, PCR on the MaxiBlock aims to both amplify and select the full-length 23,796 bp assembly product. Comparing the PCR-amplified products and the pre-PCR assembly reaction there is a clear depletion of DNA molecules composed of 10, 20, 30 and 40 fragments while the full-length assembly product of 50 eBlock represents 12% of the total DNA content in the mix (**Figure 5C**). Taken together these analyses indicate that our fully *in vitro* iterative method is able to faithfully construct long synthetic DNA molecules by assembling commercially available short dsDNA.

The cost of construction of the 23,796 bp molecule including DNA and enzymatic reactions, is about $2,000 without negotiation on suppliers’ list prices (see **Table 1**). One important aspect of our protocol is the relatively short hands-on time : once the materials have been received and the DNA sequences designed, the full-size construction process can be completed within 3-4 days. On day one, eBlocks are controlled by agarose gel electrophoresis and quantified. The assembly of the eBlocks into BigBlocks is carried out overnight. On day two, the assembled BigBlocks are controlled by agarose gel electrophoresis, followed by PCR selection. PCR products are then controlled by agarose gel electrophoresis and quantified by spectrometry. The MaxiBlock assembly is then realized overnight. On day three, the MaxiBlock assembly is controlled on a TapeStation and PCR amplified. The final PCR products are ultimately controlled on TapeStation. Higher throughput is easily achievable by robotizing and parallelizing the assembly ^25^

The length of the final DNA is limited by the capabilities of the DNA polymerase used for the final PCR selection. In fact, obtaining PCR products larger than 24 kb can be challenging with respect to efficiency and accuracy with standard DNA polymerase such as Taq polymerase. However, some DNA polymerases are known to have a higher processivity, extension rate, or error correction capabilities making them more suitable for amplifying longer DNA fragments. LongAmp^®^ Hot Start Taq DNA polymerase from NEB^®^ has the theoretical capacity to amplify DNA fragments up to 35 kb in length, while Platinum™ SuperFi II DNA Polymerase from Invitrogen™ can amplify up to 40 kb and TaKaRa LA Taq® DNA Polymerase from Takara Bio can amplify up to 48 kb. However, it is important to keep in mind that amplifying long DNA fragments can be a difficult task, and successfully obtaining a PCR product larger than 24 kb will depend on various factors such as DNA quality, reaction conditions, and the optimization of PCR conditions.

The final PCR product can be either directly used for molecular biology processes such as *in vitro* transcription with T7 RNA polymerase if a T7 promoter was included in the PCR primers. It must be highlighted that this product is not clonal and therefore contain a population of different molecules with potentially point substitutions or deletions. To obtain a clonal product, the DNA molecule produced may be purified by performing agarose gel electrophoresis and excising the appropriate size band, and cloned into a suitable host/vector system for amplification and selection. The size of the final product (∼ 24 kb) remains compatible with cloning into plasmid or cosmid vectors. The *in vitro* process described here allows for the time-efficient construction of long dsDNA molecules. The absence of any subcloning, transformation and selection step allows to streamline the process of long dsDNA molecules construction. Yet, they can be readily subcloned and selected on the basis of a double PCR screen using 2 pairs of primers located in the vector backbone and the building blocks at the extremity. Therefore, we feel this protocol may be an approach of choice to obtain artificial building blocks of thousands of nucleotides long that can be used in downstream applications.

## Methods

### Encoding of a binary file to DNA alphabet

We encoded the first articles of the Declaration of Human and Citizen Rights (DDHC) as a binary string of 4.2 Ko (**Supp data 1**). The binaries were first randomized by performing bitwise XOR operation between the input binary and the output of the hash function (SHA-256) over a constant in a process similar to what is described in (^26^). The randomized binary is converted to DNA alphabet using an algorithm that respects synthesis rules of the DNA blocks, briefly, 60 > %GC > 40, no homopolymer > 3 nt (see **Supp data 5**). This encoding allows to alter the DNA sequence at the β-encoded bits to ensure the absence of *BsaI* restriction sites and of inverse repeat regions longer than 10 nt. The 23,400 nt long DNA sequence (**Figure 1C and Supp data 1**) is partitioned into forty blocks of 472 nt (eight internals eBlocks for each of the five BigBlocks) and ten blocks of 452 nt (one eBlock for each BigBlock extremity). Each internal block is then framed by a DNA sequence composed of a 15 nt buffer sequence and a specific *BsaI* restriction site. Similarly, each external block is framed by a DNA sequence that includes a 20 nt region following the *BsaI* restriction site, which serves as a target for PCR selection of the BigBlock. Digestion with *BsaI* generates a 5’ overhang of 4 nt which allows ordered assembly. Overhangs are selected with the NEBridge GetSet™ Tool. Overhangs used are CGCT, ACGA, GCAA, CACC, CCTA, CGGA, TGAA, ACTC, ACAT. Primers used for amplification of assembly products were defined with ITHOS, a submodule of Genofrag(^27^, with parameters defined in **Supp data 6**). A 20 nt flanking sequence composed of a *BsaI* site and a 9 nt external buffer sequence was added to each primer to allow for the assembly of BigBlocks to MaxiBlocks (primer sequences are available in **Supp data 7**). The sequence was controlled to comply to IDT™ eBlock synthesis rules.

### DNA fragments

Fifty DNA blocks were ordered as 524 bp long eBlocks™ from IDT™ (Coralville, IA). The eBlocks™ were delivered in a 96 well plate at 10 nmol/μL in TE buffer pH 8.0 (10 mM Tris-HCl/0.1 mM EDTA). Each eBlock™ was quantified with AccuGreen™ High Sensitivity dsDNA Quantitation Kit on a Qubit® Fluorometer (Invitrogen™ Q32851), aliquoted and stored at -20°C. The eBlock™ integrities were individually verified by electrophoretic analysis on 3% agarose-TBE gels stained with GelRed^®^ (Biotium).

### BigBlock assemblies

Each of the 5 BigBlocks is composed of 10 oriented eBlocks™ described above. The assembly reaction is conducted in a 25.0 μL assembly reaction mix composed of 1.0 μL NEBridge^®^ Golden Gate Assembly Kit *BsaI*-HF^®^ v2 (NEB^®^ E1601), 2.5 μL T4 DNA ligase buffer 10X (NEB^®^ B0202), 0.04 pmol of each eBlock™. Reactions were conducted for 65 cycles (37°C, 5’ ; 16°C, 5’) and stopped by incubation at 60°C for 5’. The expected assembly size is 4,764 bp.

BigBlocks were PCR-amplified with dedicated primer pairs in a 25.0 μL PCR reaction mix composed of 0.25 μL Q5^®^ Hot Start High-Fidelity DNA Polymerase 2 U/μL (NEB^®^ M0493), 5.0 μL Q5^®^ 5X buffer (NEB^®^ B9027), 0.75 μL dNTP 10 mM Mix (NEB^®^ N0447), 1.25 μL of each primer (10 μM) and 1.5 ng of BigBlock as template. Amplification was conducted at 98°, 30’’ ; 5x (98°C, 10” ; *Thyb*, 20” ; 72°C, 2’30) followed by 10x (98°C, 10” ; 72°C, 2’30) and a terminal extension at 72°C, 2’00. The annealing temperature (*Thyb*) was 62°C except for BigBlock 1, 66°C and BigBlock 3, 66°C. The expected size of each amplified BigBlock is 4,796 bp. BigBlock assemblies and PCR products were analyzed by electrophoresis on 1-2% agarose-TBE gels stained with GelRed^®^ (Biotium).

### MaxiBlock assembly

The MaxiBlock is assembled from the five BigBlocks (BigBlock 1 to 5) in a 20.0 μL reaction mix composed of 2.0 μL NEBridge^®^ Golden Gate Assembly Kit *BsaI*-HF^®^ v2 (NEB^®^ E1601), 2.0 μL T4 DNA ligase buffer 10X (NEB^®^ B0202) and 200 ng of each BigBlock. Reaction was conducted for 65x (37°C, 5’ ; 16°C, 5’) and stopped by incubation at 60°C for 5’. The MaxiBlock size is 23,796bp.

The MaxiBlock was PCR-amplified with primers MaxiBlock_PCR_Fw and MaxiBlock_PCR_Rv in a 25.0 μl reaction composed of 1.0 μL LongAmp® Hot Start Taq DNA polymerase, 2.5 U/μL (NEB^®^ M0534), 5.0 μL LongAmp^®^ Taq 5X buffer (NEB^®^ B0323), 0.75 μL dNTP 10mM Mix (NEB^®^ N0447), 0.25 μL primers pair 5,0 μM and 1.0 ng MaxiBlock assembly. Amplification was conducted at 94°, 30’’ ; 15x (48°C, 20” ; 55°C, 20” ; 65°C, 22’) and a terminal extension at 65°C, 15’. The expected size of each amplified MaxiBlock is 23,796 bp.

MaxiBlock assembly and PCR products were analyzed on TapeStation 4200 (Agilent-Technologies) following the supplier protocol using Genomic DNA reagents (Agilent-Technologies 5067-5366) loaded onto a Genomic DNA ScreenTape (Agilent-Technologies 5067-5365).

### DNA sequencing

BigBlock and MaxiBlock assemblies and PCR products were sequenced using the Oxford Nanopore Technologies (ONT) MinION Mk1C and GridION Mk1 sequencer hardware respectively. The sequencing libraries were prepared according to the manufacturer protocol (Version: NBE_9065_v109_revAD_14Aug2019) with Ligation Sequencing Kit (ONT SQK-LSK109) and PCR-free Multiplexing Native Barcoding Kit (ONT EXP-NBD104). Sequencing libraries were quantified with AccuGreen™ High Sensitivity dsDNA Quantitation Kit on a Qubit^®^ Fluorometer (Invitrogen™ Q32851) and loading on Flowcell Flongle (ONT FLO-FLG001) for a run time of 24 hours.

### Data processing

Raw sequencing data were preprocessed with ONT Guppy software (V6.0.1 and V6.3.9, ^28^) for super accuracy base calling (see config file **Supp data 8**) and demultiplexing (front and rear score set > 60). Sequencing quality of each demultiplexed samples were controlled with Nanoplot (V1.38.0, ^29^). Scripts are available as **Supp data 9. S**equencing data are made available on the European Nucleotide Archive under accession number **PRJEB62556**.

### Bioinformatics results analysis

Each quality passed long read is analysed to identify the order and the identity of the blocks that compose it. We used the Smith-Waterman algorithm (^30^) to test the local alignment of a read to all the 50 original block sequences. The highest scoring block is identified and removed from the read. The alignment procedure is then recursively applied to the other parts of the read before and after the aligned sub-sequence of the read, until the entire read is identified in blocks or not. To quantify the association between each block, we determined whether each individual block is followed by an identified block, an unrecognized block or the end of the read. To analyse the global assembly, we classified reads as either *correct* if all the blocks composing it are correctly ordered, or as *incorrect* if at least one block is misplaced.

## Supporting information

Time table

quality metrics

short sequence analysis

encoding text into DNA

primer selection parameters

primer sequences

configuration file

scripts

## Author Contribution

JL, ER, DL, JN, OB, YA Olivier Boulle *Investigation, Formal analysis*

DL *Supervision, Funding acquisition, Project administration, Conceptualization*

JN *Conceptualization, review and editing*

JL *Investigation, Conceptualization, review and editing*

ER *Conceptualization, review and editing*

YA *original draft preparation, review and editing, Supervision, Conceptualization*

## Acknowledgment

Julien Leblanc is supported by the Labex CominLabs (dnarXiv project), Olivier Boulle is supported by the PEPR MolecularXiv (ANR-22-PEXM-003). We gratefully acknowledge Edouard Cadieu and Thomas Derrien (IGDR) for their help with ONT sequencing and data processing.

## Supporting Information

Supplementary data 1 - Text and DNA encoding of the Universal Declaration of Human rights and citizens, sequences of the ordered geneblocks. (TXT)

Supplementary data 2 - supplementary figure presenting the time table of the assembly of a 24 Kb artificial dsDNA fragment. (PDF)

Supplementary data 3 - ONT sequencing quality metrics. (TXT)

Supplementary data 4 – Supplementary figure presenting the analysis of unidentified reads. (PDF)

Supplementary data 5 - Supplementary figure presenting the DNA encoding scheme of a text file.

(PDF)

Supplementary data 6 - primer selection parameters. (TXT)

Supplementary data 7 – oligonucleotides sequences. (TXT)

Supplementary data 8 - configuration file for ONT Guppy basecaller software (TXT)

Supplementary data 9 – Script file for basecalling, demultiplexing and quality control

