## Supplementary material for "“In vitro construction and long read sequencing analysis of a 24 kb long artificial DNA sequence encoding the Universal Declaration of the Rights of Man and of the Citizen ”": Time table

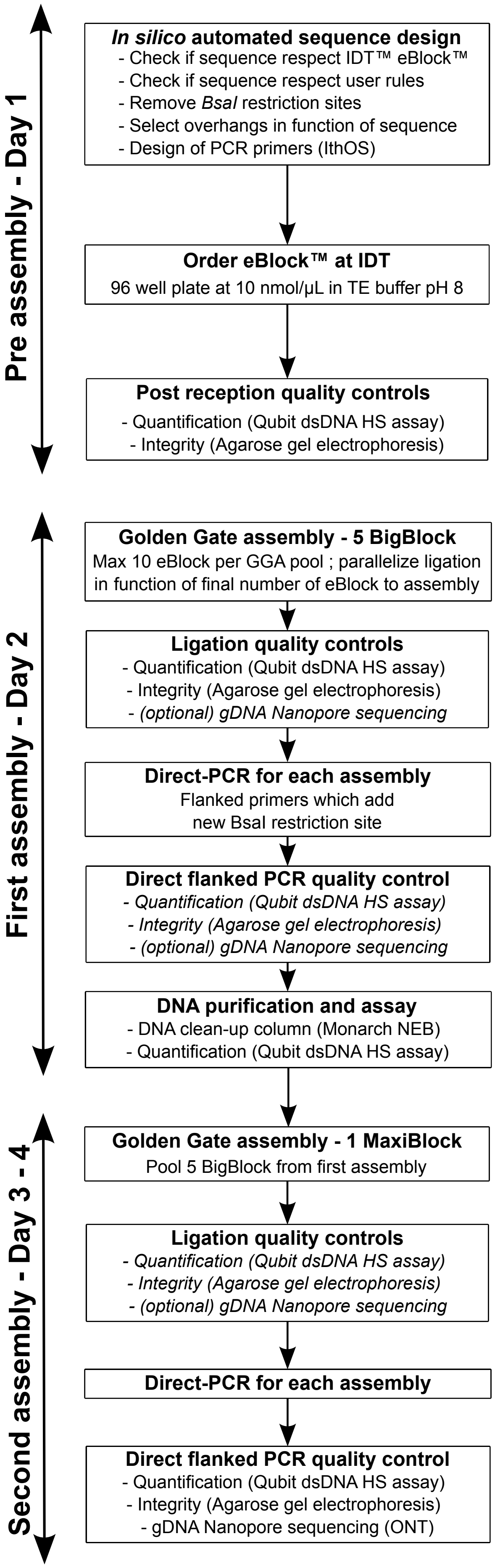


**Supplementary data 2: Time table of the assembly of a 24 Kb artificial dsDNA fragment**

Pre-assembly - 1 day plus eblock production. Starting from a desired sequence, a program generates the sequences of the different parts (eBlocks) composing the final DNA molecule in compliance with the IDT constraints for eBlock synthesis. Overhangs used for GGA are chosen according to the NEBridge™ Ligase Fidelity web tool. eBlocks are ordered. Flanked primer targeting the different BigBlocks are designed for the PCR selection process.Upon arrival, the eBlocks are quantified using Qubit HS assay and their integrity verified by agarose-gel electrophoresis.

First-assembly - Day 2. Five BigBlock of 5Kb are build in vitro. Five equimolar pools of 10 eBlocks are assembled by GGA into BigBlocks in parallel and checked by agarose-gel electrophoresis before proceeding to the PCR selection with flanked primers. To measure the efficiency of the ordered assemblies, ligation products and PCR selected ligation products are controlled by sequencing on a MinION Mk1C sequencer. PCR amplified BigBlocks are column purified and quantified before proceeding to the second assembly step..

Second-assembly - Day 3 and 4. The five BigBlock of 5Kb are assembled into one MaxiBlock of ~24 Kb by GGA. The expected full-length product (MaxiBlock) is specifically amplified using primers hybridizing to the extremity of first and last eBlocks composing the DNA molecules. Both ligation products and PCR amplified DNA have been controlled using capillary electrophoresis on a Genomic DNA ScreenTape System because of the long size of the DNA molecule and by sequencing on GridION X5 Mk1 sequencers.
