## Supplementary material for "“In vitro construction and long read sequencing analysis of a 24 kb long artificial DNA sequence encoding the Universal Declaration of the Rights of Man and of the Citizen ”": short sequence analysis

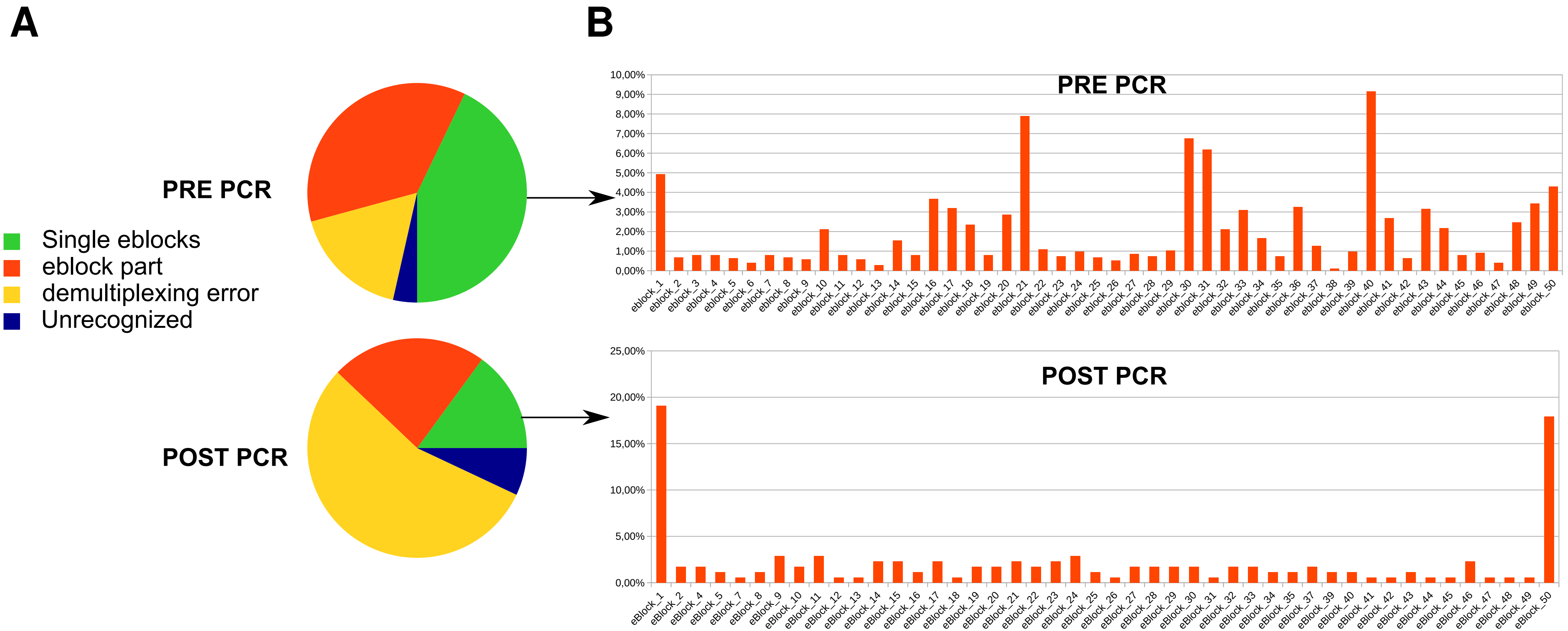


**Supplementary data 4**: Analysis of short sequences present in the MAXiblock sequencing experiment.
**A)** Short sequences present in the pre or post PCR fraction are composed of 1) single eBlock fragments, 2) parts of eblocks, 3) oligo sequence from a distinct experiment (demultiplexing error) or truly unrecognized fragments. **B)** Short sequences obtained before and after PCR AND containing eblock sequences are mainly composed of single eblocks with eblocks 1 and 50 being enriched in the post-PCR fraction.
