## Supplementary material for "“In vitro construction and long read sequencing analysis of a 24 kb long artificial DNA sequence encoding the Universal Declaration of the Rights of Man and of the Citizen ”": encoding text into DNA

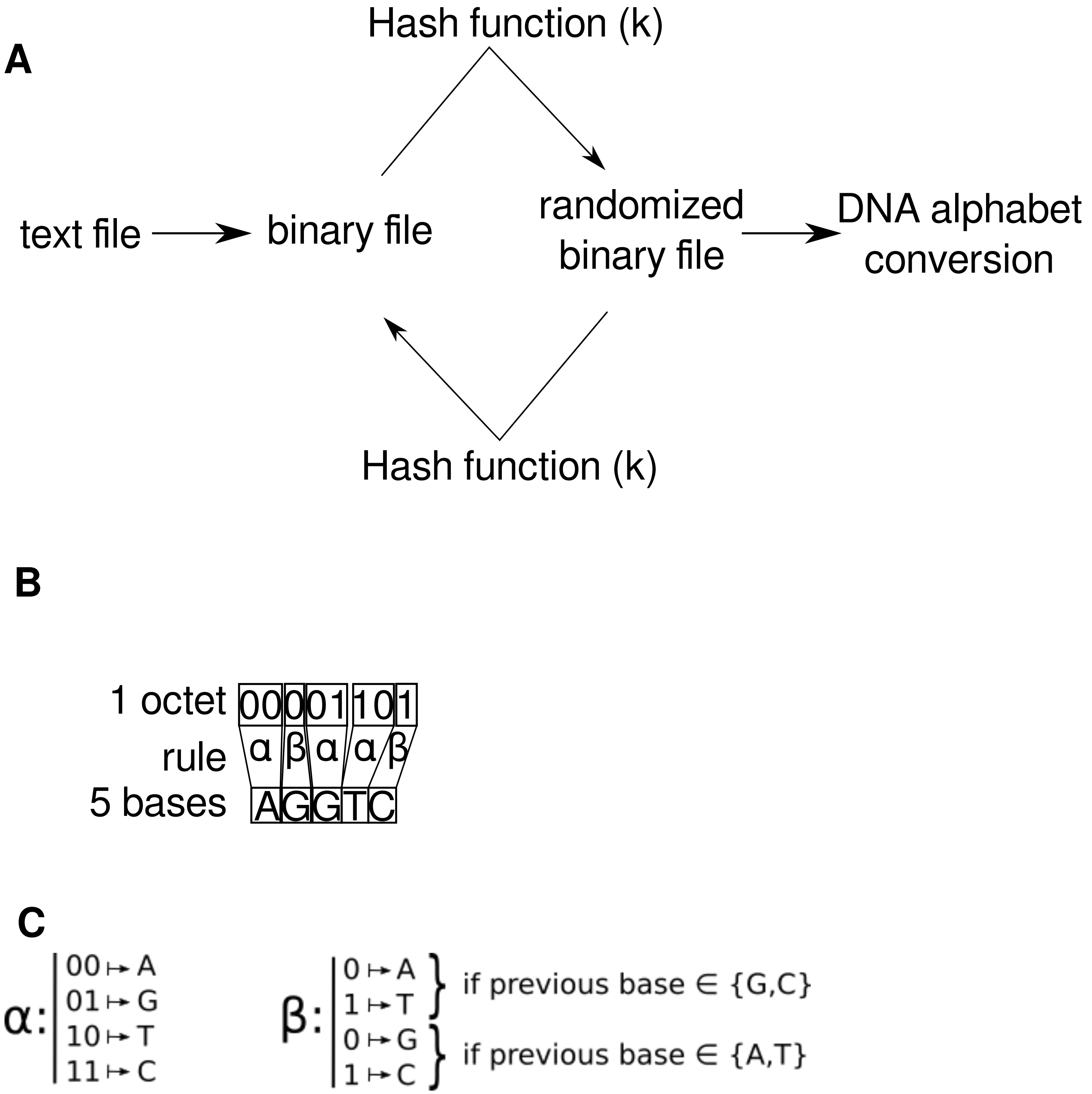


**Supplementary data 5**: encoding of a binary file in a DNA alphabet.
a) the text file is a binary file. It is randomized using a bitwise XOR operation of the binary file and the output of a hash function. The same hash function allow to restore the original binary file from the randomized one. The randomized file is converted to a DNA alphabet using the set of rule defined in B and C. B) Each bit of the randomized binary file is converted to a DNA base alphabet following rule α (2 bits/nt) or β (1 bit/nt) following the pattern αβααβ. C) Definition of rule α and β.
Since a β-coded base is always different from the preceding α-coded base, it breaks any possible homopolymers > 3; and it also rebalances the %GC content of the quintuplet of bases to 40-60%.
The total encoding rate is 1.6 bits/nt ( [3*2 bits/nt + 2*1bit/nt] /5 ).
